# Unlocking cellular plasticity: Enhancing human iPSC reprogramming through bromodomain inhibition and extracellular matrix gene expression regulation

**DOI:** 10.1101/2023.10.13.562265

**Authors:** Jun Yang, H. Karimi Kinyamu, James M. Ward, Erica Scappini, Trevor K. Archer

## Abstract

The transformation of fibroblasts into epithelial cells is critical for iPSC reprogramming. In this report, we describe studies with PFI-3, a small molecule inhibitor that specifically targets the bromodomains of SMARCA2/4 and PBRM1 subunit of SWI/SNF complex, as an enhancer of iPSC reprogramming efficiency. Our findings revealed that PFI-3 induces cellular plasticity in multiple human dermal fibroblasts, leading to a mesenchymal-epithelial transition (MET) during iPSC formation. This transition was characterized by the upregulation of E-cadherin expression, a key protein involved in epithelial cell adhesion. Additionally, we identified COL11A1 as a reprogramming barrier and demonstrated COL11A1 knockdown increased reprogramming efficiency. Notably, we found that PFI-3 significantly reduced the expression of numerous extracellular matrix (ECM) genes, particularly those involved in collagen assembly. Our research provides key insights into the early stages of iPSC reprogramming, highlighting the crucial role of ECM changes and cellular plasticity in this process.

## INTRODUCTION

Cellular plasticity is the ability of cells to transform from one fate to another [1]. It is a crucial process in various cellular functions, including cell development, wound repair, and cancer progression [2]. Epithelial to mesenchymal transition (EMT) exemplifies cellular plasticity in cancer cells. During iPSC reprogramming, fibroblasts, of mesenchymal origin, are initially transformed into epithelial cells through a process known as mesenchymal to epithelial transition (MET) [3]. MET is an essential step in cell reprogramming since iPSCs exhibit characteristics similar to epithelial cells, such as tight junctions and polarity [4]. The plasticity of fibroblasts, as demonstrated by MET during iPSC reprogramming, is closely associated with the efficiency of reprogramming [5]. As such, this cell type switch involves extensive changes in gene expression, including the activation and inhibition of genes in both cell types [6].

Cellular plasticity is regulated by various factors, including epigenetics, chromatin modifiers [5], and extracellular matrix (ECM) [1]. Bromodomains (BRD) are protein domains that recognize and bind acetylated lysine on histones [7]. These BRD proteins, classified as epigenetic readers, orchestrate gene expression via mechanisms such as chromatin remodeling, histone recognition, and histone acetylation [8] [9]. PFI-3, a bromodomain inhibitor, specifically targets SMARCA2/4 (BRG1/BRM) and PBRM1 (BAF180) subunits of SWI/SNF chromatin remodeling complex [10, 11]. Recent studies have shown that PFI-3 inhibits the binding of BRG1 to chromatin and displaces BRG1, BRM, and BAF180 from chromatin, depending on cellular context [12]. However, the specific role of these chromatin remodeling enzymes in cell plasticity is not fully understood. Previous research has shown that BRG1 regulates gene expression related to extracellular matrix (ECM) and adhesion in melanoma cells [13], as well as genes involved in cell polarity and cell adhesion in retinal development [14]. Similarly, BAF180 has been found to regulate expression of genes related to cell adhesion in renal carcinoma [15]. These studies indicate that chromatin remodeling enzymes collaborate with ECM to regulate gene expression and cell plasticity [16].

The ECM is a dynamic protein network [17] that regulates biochemical and mechanical signals essential for determining cell fate, including differentiation, adhesion, and migration [1, 18]. The ECM constantly undergoes remodeling process such as assembly and degradation [19]. Remarkably, changes in collagen deposition and collagen crosslinking alter the stiffness of ECM [20, 21], which in turn promotes cell migration, invasion, and EMT through mechanotransduction [22]. EMT in cancer metastasis has been extensively studied [23, 24], while the reverse process, MET, and its underlying mechanisms are not well understood. Interestingly, both carcinogenesis and iPSC reprogramming processes have similarities [25]. During iPSC reprogramming, MET is an essential step before establishing pluripotency [26]. However, the induction of MET during reprogramming has yet to be fully elucidated.

In this study, we found that bromodomain inhibitor PFI-3 increased reprogramming efficiency in human dermal fibroblasts. Further experiments revealed that PFI-3 altered the gene expression of ECM and promoted MET during iPSC reprogramming. These findings provide valuable insights into the mechanisms underlying cellular plasticity and the regulation of gene expression by the ECM. Understanding these processes could potentially lead to improved iPSC reprogramming techniques and advancement in regenerative medicine.

## MATERIALS AND METHODS

### Cell lines and Cell culture

The establishment of primary dermal fibroblast was carried out according to the methods in our previous published work [27]. In summary, fibroblast lines were generated from skin biopsies acquired from NIEHS Clinical Research Unit. Skin punch biopsies, each having a diameter of 4mm, were cut into small pieces, and dried on 60mm petri dish for 5 minutes and then added DMEM (Gibco 11965-092) with 10% FBS (Atlanta Biologicals) and 1% Pen/Strep (Sigma). Media was changed daily until fibroblasts emerge.

### Reprogramming Assay

iPSC generation was conducted as described in our previous study [27]. Briefly, fibroblasts were reprogrammed using lentiviral vectors that carried six transcription factors (ADDGENE/PSIN4-EF2-N2L, ADDGENE/PSIN4-EF2-O2S and ADDGENE/PSIN4-CMV-K2M). Fibroblast cells were plated in 6-well plates at 1.5 x 10^5^ cells per well (Day 0). On Day 1, these cells were transduced using viral soups (titer of 6×10^6)^ and 8ug/ml of polybrene (Day 1). Virus was removed after 24 hours and replaced with fresh DMEM complete media for 48 hours. Cells were then transferred to 10cm dishes coated with Matrigel (Corning Cat# 354234 diluted in DMEM/F-12, Gibco). 48 hours later cells were fed with a 1:1 mixture of complete DMEM and E8 media (TeSR^TM^-E8^TM^ Stem Cell Technologies). Cells were subsequently maintained in E8 daily for 21 days. PFI-3 was purchased from Sigma (SML0939) and was applied during different timeframes. The iPSC colonies were picked using a 20 uL pipet and grown up individually in E8 media. ReLeSR (Stem Cell Technologies) was used for harvesting and splitting.

Reprogramming efficiency was determined by alkaline phosphatase staining (Stemgent AP Staining Kit II). Triplicate reprogrammed dishes (10 cm) containing colonies were AP stained and scanned to images. The software program ImageJ (Wayne Rasband National Institutes of Health, USA) was used for counting colonies from color threshold adjusted and binary converted images of each dish. Triplicate plates were averaged and reported as colony counts or percent reprogramming efficiency ((# of colonies/150,000) x100).

### Embryoid Body (EB) Formation and Differentiation

iPS colonies were detached with ReLeSR and aggregated in a 6-well low attachment plate containing E8 media for 48 hours to form EBs. Subsequently, the EBs were separated from loose cells using a reversible strainer (Stem Cell Technologies Cat# 27259) and transferred to a new ultralow attachment plate with EB media (Stem Cell Technologies Cat# 05893). The media was then replaced by half daily. On day 7, the EBs were transferred to Matrigel coded 6-well plate and cultured in EB medium for 12 days, during which they will differentiate into three germ layers.

### Quantitative RT-PCR

Total RNA was extracted using RNeasy Mini Kits (Qiagen). Two microgram of RNA was reverse transcribed using the SuperScript First Strand kits (Invitrogen), following treatment with DNase I (Invitrogen). The qRT-PCR analysis was carried out using the Brilliant III Ultra-Fast SYBR Green QPCR Master Mix (Agilent Technologies) with 40 cycles of 20s at 95°C and 20s at 60°C. The housekeeping gene, TBP, was used to calculate relative expression. RT-PCR Primer sets for 15 genes which are listed in Supplement Figure 1.

### Western Blot Analysis

Whole cell lysate was isolated using Buffer X (100 mM Tris-HCL pH 8.5, 250 mM NaCl, 1% (v/v) NP-40, 1 mM EDTA) combined with protease inhibitors, phosphatase inhibitor, and PMSF (Sigma). Protein concentrations were determined using the colorimetric Bradford Protein Analysis method with BioRad reagents. Western blotting was performed on precast 18-well 7.5% Tris-HCL gels (Criterion cat# 345-0006) loaded with 50 ug of whole lysate. The gel was then transferred to nitrocellulose membranes using Tris/Glycine buffer containing 20% methanol. Membranes blotting was carried out in 5% nonfat milk for an hour, followed by overnight incubations of primary antibody. The antibodies used were as follows: COL11A1 (Sigma, HPA058335), E-Cadherin (Cell Signaling, 24E10, #3195), FN1(Sigma, F3648), LIN28 (PTG, 11724-1-AP), NANOG (Bethyl, A300-398A), OCT4 (Santa Cruz, sc-5279), SOX2 (Santa Cruz, sc-20088), β-Actin AC-15 (Sigma cat# A1978) and GAPDH FL-335 (Santa Cruz sc-25778). After incubation with appropriate secondary antibody, blots were quantified using digital fluorescence with LI-COR imaging reagents and the Odyssey™ CLX Imaging System.

### Immunostaining

Cells were fixed using 4% paraformaldehyde for 20 minutes at room temperature, then permeabilized with 0.5% Triton-X100 for 10 minutes. This was followed by blocking with 0.1% Triton-X100, 5% normal donkey serum in PBS for 1 hour at room temperature. After washing three times with PBS, cells were incubated overnight with the primary antibody at 4°C. The antibodies used were as follows: AFP (Millipore 2004189), α-SMA (Millipore CBL171), E-Cadherin (Cell Signaling, 24E10, #3195), LIN28 (PTG, 11724-1-AP), NANOG (Chemicon AB5731), Nestin (Millipore ABD69), OCT4 (Santa Cruz, sc-5279), SOX2 (Chemicon AB5603), TRA-1-60 (Millipore MAB4360), TRA-1-81 (Millipore MAB4381).

### LOX Activity Assay

Extracellular LOX enzymatic activity was quantified using Abcam LOX activity assay kit (ab112139), which measures the hydrogen peroxide generated by LOX through a proprietary red fluorescence substrate for HRP-coupled reactions. A mixture of 50 uL of cell culture supernatant and 50 uL of reaction buffer was incubated at 37°C for 10 minutes, protected from light. Assays were performed using black 96-well plates (Porvair, Cat# 205003). Fluorescence was then detected at Ex/Em = 540/590 nm using a fluorescence microplate reader. As the assay is semi-quantitative and does not contain a LOX standard, LOX activity is expressed as absolute fluorescence value.

### Knockdown of Gene Expression by shRNA

COL11A1 shRNA plasmid was obtained from NIEHS Genomic core (Thermo Scientific GIPZ Lentiviral shRNA). Lentiviral packaging was produced at the NIEHS Viral Vector Core, following a previously established protocol [28]. The successful knockdown of COL11A1 was subsequently verified by qRT-PCR.

### Gene Expression Profiling and Analysis

Total RNA was isolated using Qiagen RNeasy kit and quantified using Agilent Nanodrop. RNA quality was assessed using a 2100 Bioanalyzer instrument and an Agilent 6000 RNA Pico Kit (Agilent Technologies).

Affymetrix Clariom D human transcriptome array was processed using oligo (https://doi.org/10.1093/bioinformatics/btq431) and analyzed using limma (https://doi.org/10.1093/nar/gkv007) R packages. Probe-level statistical results were combined to genes by Fisher’s method, P-values were corrected by Benjamini Hochberg (ref https://doi.org/10.1111/j.2517-6161.1995.tb02031.x). Significant genes had adjusted P-value 0.05 and fold change 1.25.

Analyses were performed independently within batch, then statistical results combined for downstream visualization.

### Gene Ontology and Pathway Analysis

Pathway analyses used clusterProfiler (https://doi.org/10.1089/omi.2011.0118) with significant genes across days 3, 5, and 7, versus Hallmark and canonical pathways from MSigDB (https://doi.org/10.1093/bioinformatics/btr260). Significant pathways had an adjusted P-value of 0.05 and at least four genes.

Violin plots show mean abundance of significant pathway genes by condition, and lines indicate collective changes in expression, with paired t-test P-value displayed above each comparison.

### Pathway Enrichment Analysis

Ingenuity pathway analysis (IPA; Qiagen), was run using default settings for genes with Spearman correlations with p ≤ 0.01 (0.388 for n=36 and 0.274 for n=72) for the total cohort combined and ancestries separately.

Enriched pathways with an FDR adjusted p-value less than 0.1 were used to generate a pathway-gene heatmap, which was clustered using Euclidian distance. Concept network (Cnet) plots were created using exemplar pathways from each cluster.

### Enrichment map network

Enrichment Map networks were created using pathways with an adjusted P-value of 0.05, where pathways were colored by direction of gene regulation, and were connected between pathways with a Jaccard gene overlap at least of 0.2.

## RESULTS

### Bromodomain inhibitor PFI-3 enhances reprogramming efficiency of human dermal fibroblasts

Small molecule PFI-3 specifically targets SMARCA2/4 (BRG1/BRM) and PBRM1 (BAF180) subunit of SWI/SNF complex [10, 11]. Given the important roles of bromodomain proteins in transcriptional regulation and biological processes, we investigated whether PFI-3 could enhance iPSC generation. To determine the effect of PFI-3, human dermal fibroblasts (African American Female) were reprogrammed using Yamanaka factors OCT4, SOX2, KLF4, c-MYC, LIN28, and NANOG (OSKMLN) and treated with PFI-3 for varying time periods (Figure 1A). Treatment of PFI-3 for the entire 21-day reprogramming process led to a three-fold increase in iPSC colony formation (Figure 1B). However, pretreatment of fibroblasts with PFI-3 for 10 days prior to reprogramming had no effect on reprogramming efficiency (Figure S1A). To validate this finding, multiple fibroblast cell lines from different ancestry and gender were tested, and it was found that PFI-3 consistently improved reprogramming efficiency in human dermal fibroblasts (Figure S1B).

**Figure 1.**
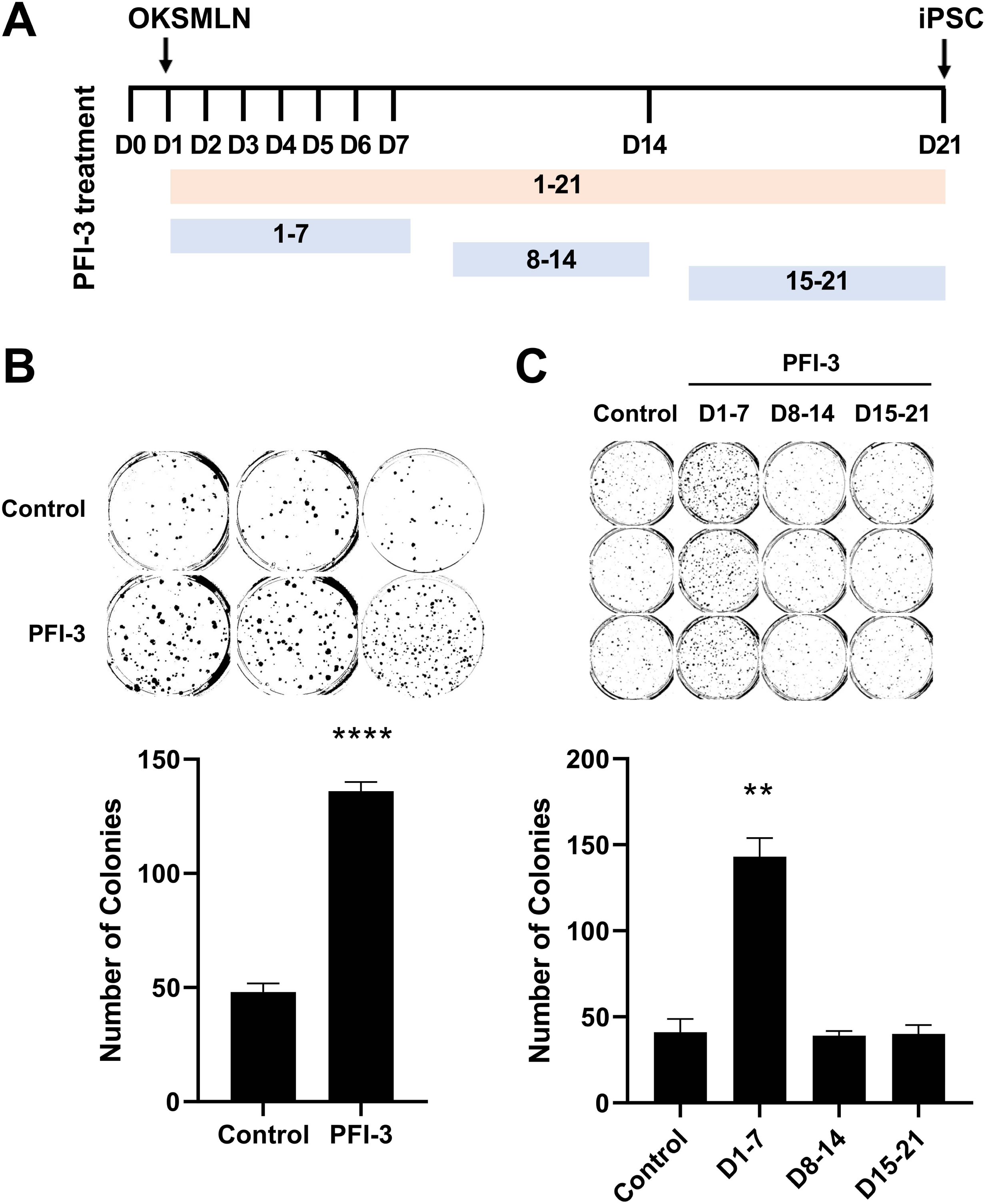
Brg-1 bromodomain inhibitor PFI-3 enhances reprogramming efficiency of human dermal fibroblasts. **A)** Schematic protocol for PFI-3 treatment intervals during iPSC reprogramming. **B)** PFI-3 promoted human iPSC reprogramming. Human dermal fibroblast cells were transduced with OSKMLN and were treated with 10 uM PFI-3 for the entire 21 days. AP staining at day 21 shows a high reprogramming activity of PFI-3. n=3. *P* values were determined by a two-tailed Student’s *t*-test; \*\*\*\**P* value <0.0001, n=3. *P* value was 0.00006. **C)** Time course of PFI-3 reprogramming activity. PFI-3 has effect on the first 7 days. \*\**P* value <0.01, n=3. *P* value were 0.0015, 0.85, 0.89.

Further investigations were conducted to determine the optimal timing for PFI-3 treatment to enhance iPSC generation. Human dermal fibroblasts were transduced with OSKMLN and treated with PFI-3 at different time intervals (days 1-7, 8-14, 15-21). The results revealed that PFI-3 inhibition was most effective within the first seven days after OSKMLN transduction. Treatment during later time periods had no significant impact on reprogramming efficiency (Figure 1C).

Based on these findings, we concluded that PFI-3 bromodomain inhibition acts early in reprogramming process to enhance iPSC generation.

### PFI-3 iPSCs are pluripotent

We next investigated if the iPSCs generated using PFI-3 bromodomain inhibitor were pluripotent. PFI-3 iPSC clones were stained to test expression of cell surface markers TRA1-60 and TRA1-81 as well as the transcription factors OCT4, SOX2, and NANOG. The presence of these markers indicates pluripotency. As expected, all the PFI-3 iPSC clones exhibited positive expression of these pluripotency markers (Figure 2A), suggesting they that they possess the ability to differentiate into multiple cell types. To further confirm the pluripotency of the PFI-3 iPSCs, quantitative RT-PCR and Western blot analysis were conducted to analyze the RNA and protein expression levels of key pluripotency markers. The level of OCT4, LIN28, SOX2, NANOG, in PFI-3 iPSCs in the PFI-3 iPSCs were found to be comparable to those of a human embryonic stem cell (ESC) line H9 and control iPSCs (Figure 2B & Figure 2C). These findings suggests that PFI-3 iPSCs possess similar pluripotency characteristics to human ESCs.

**Figure 2.**
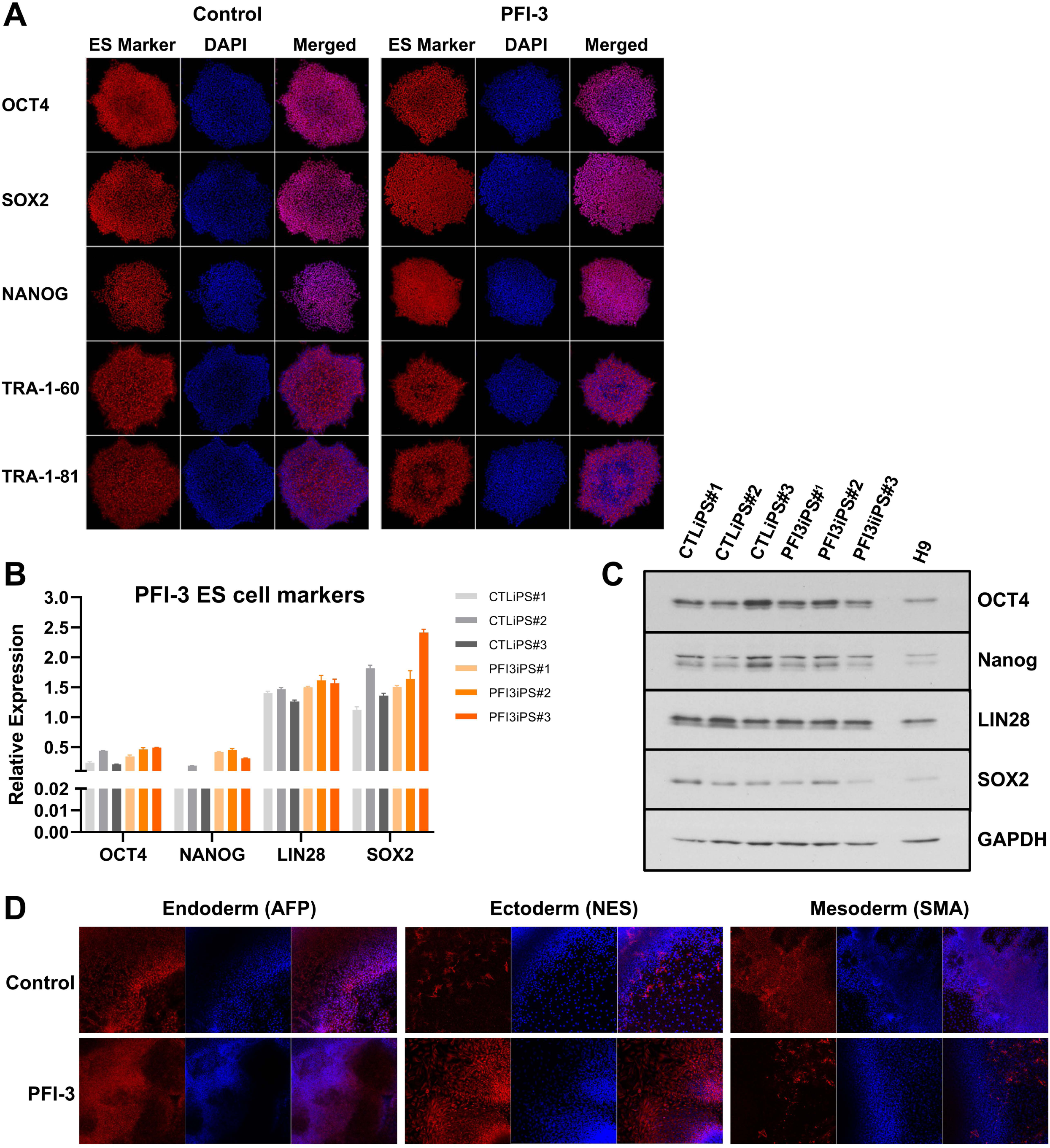
PFI-3 iPSCs are pluripotent. **A)** Immunofluorescent staining of PFI-3iPSCs shows expression of pluripotency makers OCT4, SOX2, and NANOG as well as the surface markers TRA-1-60, and TRA-1-81. **B)** mRNA expression levels of the pluripotency makers OCT4, NANOG, LIN28, and SOX2of all iPSC lines. **C)** Protein expression of the pluripotency makers of all iPSC clones. **D)** Immunofluorescent staining of PFI-3iPSCs shows expression the markers of three germ layers.

To access the developmental potential of PFI-3 iPSCs, their ability to differentiate into the three germ layers was evaluated through the formation of embryoid body (EB). The PFI-3 iPSCs were aggregated in a low attachment plate for 48 hours to form EBs. These EBs were then cultured in EB medium on Matrigel coated plates for 14 days. The resulting cells were stained positive for markers specific to endodermal, ectodermal, and mesodermal lineages, namely alpha fetoprotein, nestin, and smooth muscle actin respectively (Figure 2D). These findings further confirm the pluripotent nature of the PFI-3 iPSCs.

### Gene expression analysis of the effects of PFI-3 on iPSC reprogramming

To further examine the molecular mechanism of PFI-3 on reprogramming, we compared gene expression profiles between PFI-3 treated and control. Firstly, we performed a time course analysis at days 3, 5, and 7 of reprogramming to capture the dynamic nature of gene expression during this process. Our results showed that the gene expression profiles of PFI-3 treated sample were distinguishable from the control at each time point, indicating that PFI-3 has significant impact on gene expression during reprogramming (Figure 3A).

**Figure 3.**
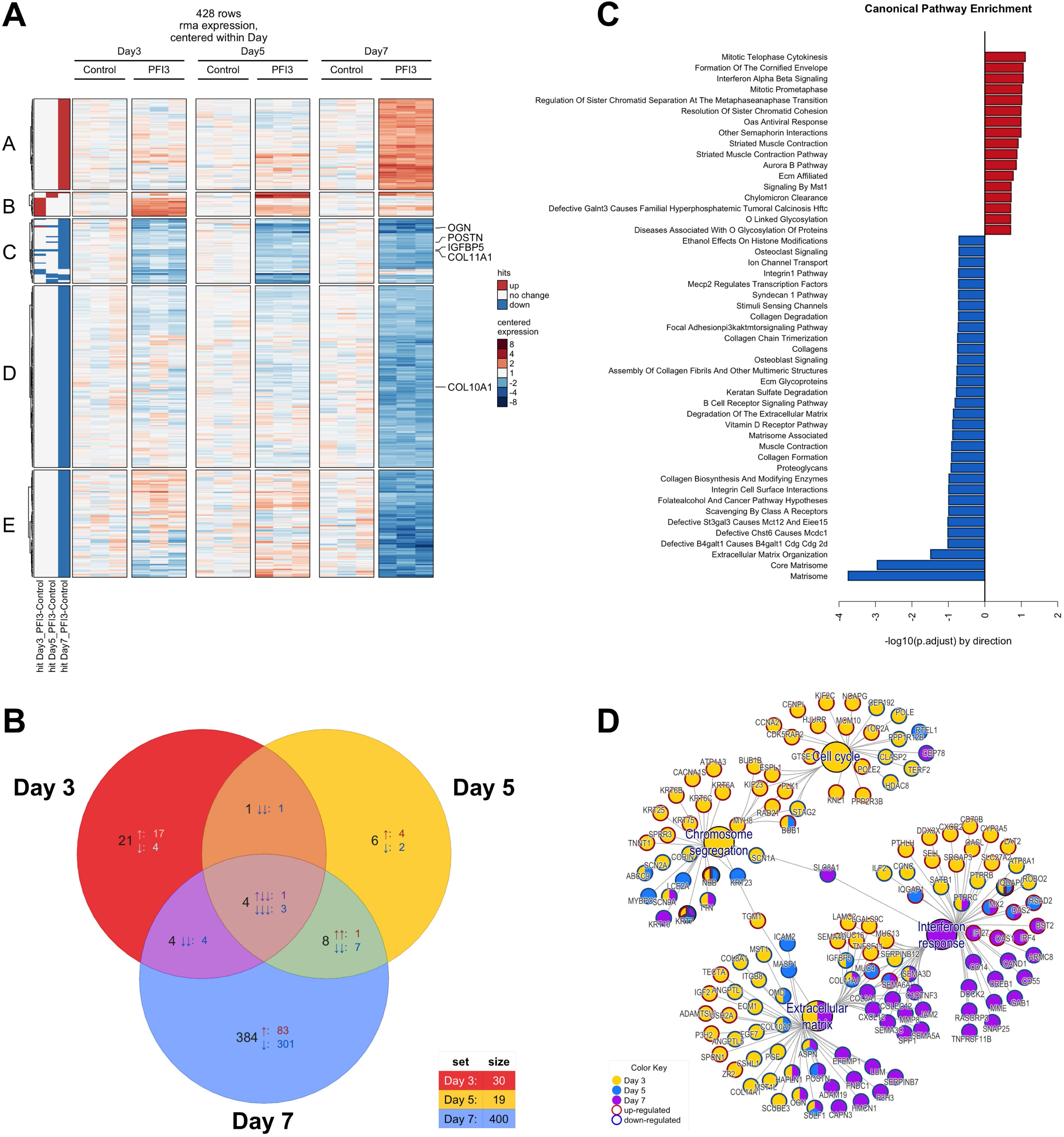
Gene expression profiling analysis. **A)** Heat map of 428 DEGs (FC > 1.5 and FDR < 0.05) in the comparison of PFI-3 and untreated across time points of reprogramming. Red denotes high and blue denotes low gene expression levels. Color scale represents the log2-fold change compared to untreated control. Genes are divided into 5 clusters according to their dynamic expression profiles**. B)** Venn diagram showing differentially common expressed genes across time points. **C)** Gene Ontology (GO) analysis of PFI-3 effects on reprogramming. Gene ontology [−log10 (p value)] biological processes of DEGs (log2FC > ±0.5 and p < 0.05) at first 7 days of reprogramming. **D)** Cnet plot of enriched pathways.

We identified a total of 428 differentially expressed genes (DEGs) between PFI-3 treated and control samples. These DEGs were clustered into five groups based on their expression patterns. The gene are listed in additional file (Table S1). Cluster A consisted of immune response related genes, which were highly enhanced by PFI-3 at day 7 (Figure S2). This suggests that PFI-3 may modulate the immune response during iPSC reprogramming. Cluster B included genes involved in cell cycle, mitosis, cytokinesis, and keratinization. These genes were upregulated by PFI-3 in the early phase of reprogramming, but their expression levels decreased over time.

Keratinization plays a key role in promoting epithelialization and safeguarding against stress-induced damage [29, 30]. Thus, it indicates that PFI-3 may promote the cell cycle progression and keratinization, ultimately facilitating the reprogramming process. Cluster C consisted of ECM glycoproteins and proteoglycans related genes. We observed that PFI-3 repressed the expression of these genes at very early time point and further downregulated them over time. ECM glycoproteins are essential components of the extracellular matrix, providing structural support and facilitating cell adhesion and migration [31]. On the other hand, proteoglycans play a vital role in cell signaling processes by regulating the activity of growth factors and cytokines, as well as controlling cell adhesion, migration, and proliferation [32]. The downregulation of these genes suggests a disruption in signal pathways essential for reprogramming [33]. Cluster D was annotated as ECM organization, focal adhesion, and collagen assembly related genes. These genes regulate cell proliferation, differentiation, and adhesion [18], PFI-3 slightly downregulated their expression at day 3 and day 5, but their expression was highly repressed at day 7. This implies that PFI-3 may impair cell-matrix interactions and remodeling processes, potentially impacting the morphological changes during reprogramming. Lastly, cluster E consisted of genes involved in receptor kinase signaling, which were highly repressed upon PFI-3 treatment at day 7. This finding suggests that PFI-3 may affect receptor mediated pathways, further highlighting its potential role in modulating cellular response during reprogramming.

PFI-3 treatment of reprogramming fibroblast resulted in extensive transcriptional changes, with most genes being repressed at day 7. Analysis of three distinct datasets revealed a total of four genes, namely ASPN, COL11A1, IGFBP5, and KRT7, that displayed differential expression. Among these differentially expressed genes, 106 genes were upregulated, and 322 genes were downregulated (Figure 3B). To gain insight into the functional roles of DEGs at each time points, we conducted Gene Ontology (GO) enrichment analysis. Our results revealed that ECM organization, collagen assembly, cell adhesion, cell cycle and cell division, and immune response were the major affected pathways (Figure 3C). The enrichment of these pathways was confirmed by Cnet plot analysis (Figure 3D).

Taken together, the gene expression analysis highlights the impact of PFI-3 on gene regulation and functional pathways involved in iPSC reprogramming process. The extensive transcriptional changes observed during the reprogramming process with PFI-3 treatment, particularly the massive gene repression on day 7, signify a critical stage in the cell transformation process. PFI-3 treatment alters the expression of genes associated with ECM organization, collagen assembly, cell-matrix interaction, and ECM remodeling, indicating a shift in cell identity and remodeling of the extracellular environment.

### PFI-3 promotes MET

Gene expression profiling indicated that PFI-3 significantly influences the gene expression associated with ECM. The ECM is a complex protein network that affects cell growth, differentiation, migration, and invasion[18]. MET is a process in which migratory cells dedifferentiate into polarized epithelial cells. This process is essential in iPSC reprogramming [26]. Interestingly, EMT, the opposite process of MET, is associated with alteration in the ECM during cancer progression. ECM has been known to drive EMT during tumor genesis. Hence, we can infer that PFI-3 might promote MET by altering the ECM composition and cell-matrix interaction, consequently changing cell morphology and plasticity.

To investigate this hypothesis, we examined the expression of the MET marker gene E-cadherin (CDH1) during the reprogramming process. We observed that E-cadherin RNA expression was upregulated in PFI-3 treated cells at day 7 (Figure 4A), while E-cadherin protein expression was detected at day 10, significantly upregulated by PFI-3 (Figure 4B). Moreover, the immunostaining demonstrated higher E-cadherin expression in in PFI-3 treated cells compared to the control (Figure 4C). Additionally, the expression of E-Cadherin correlates with the induction of pluripotent markers (Figure S3). Therefore, these findings suggest that PFI-3 promotes MET during iPSC reprogramming.

**Figure 4.**
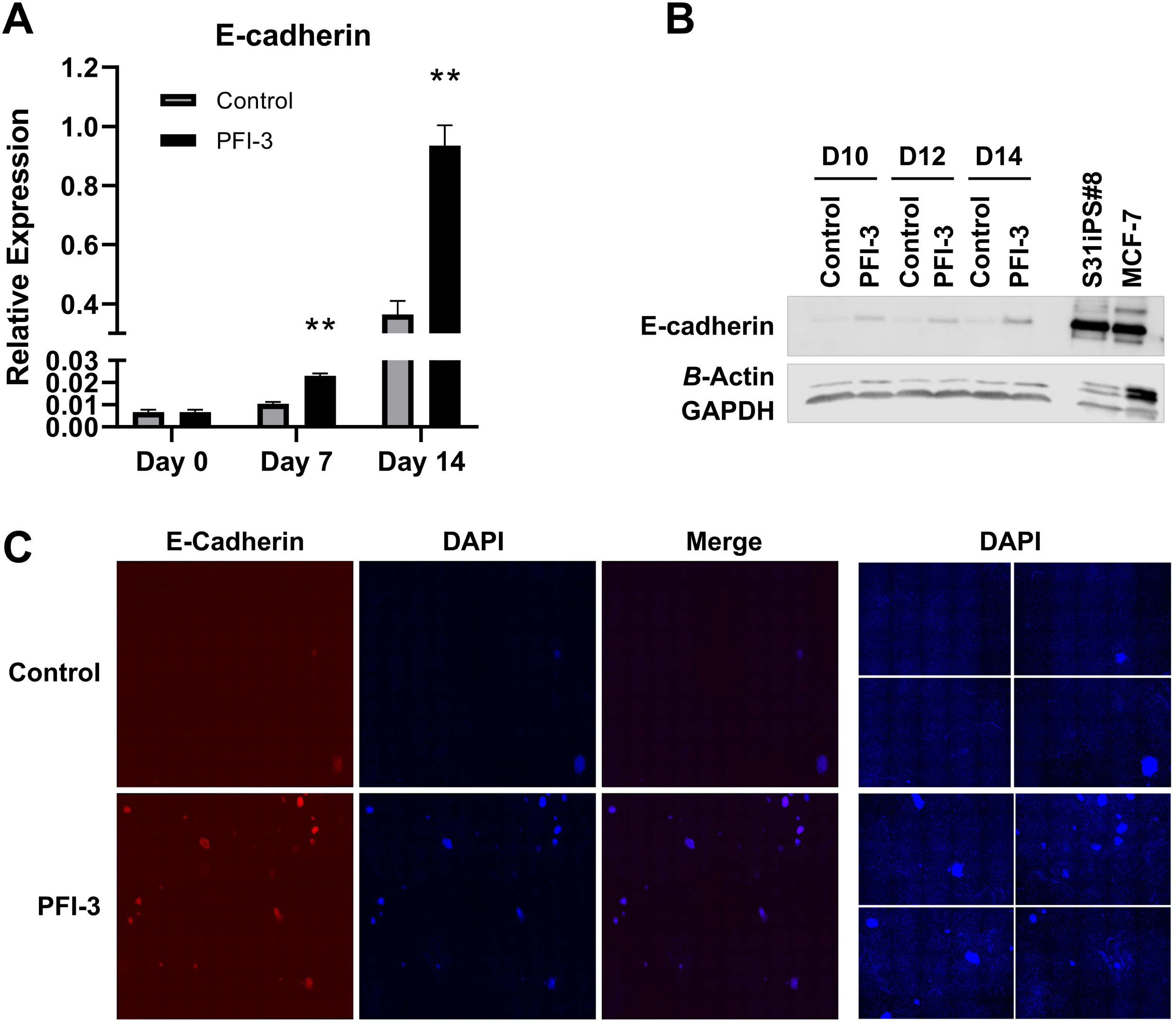
PFI-3 promotes mesenchymal to epithelial transition (MET). **A)** Relative expression level of E-cadherin (CDH1) on days 0, 7, 14 of reprogramming. Transcript level were normalized to TBP expression. \*\**P* value <0.01, n=3. *P* values were 0.0011, 0.0014. **B)** E-cadherin protein expression on days 10, 12, and 14 of reprogramming. **C)** Immunofluorescent staining of PFI-3 treated cells shows enhanced expression of MET maker E-cadherin on day 14 of reprogramming. DAPI staining shows equal cell numbers between PFI-3 treated cells and control.

### PFI-3 inhibits ECM gene expression specifically genes involved in collagen assembly and crosslinking

Gene expression profile revealed that many ECM genes were modified by PFI-3 (Figure 3A). Next, we investigated how ECM changes affected MET and reprogramming. Enrichment analysis of ECM associated gene networks revealed that genes involved in collagen assembly and focal adhesion were among the most repressed (Figure 5A). This suggested that PFI-3 may affect the structural integrity and mechanical properties of the ECM. Additionally, biological network analysis demonstrated that PFI-3 had significant effects on cell cycle and cell division, cell morphology, and mechanical stimulus (Figure 5B). This further supports the notion that PFI-3 plays a role in ECM remodeling and cellular processes related to ECM dynamics.

**Figure 5.**
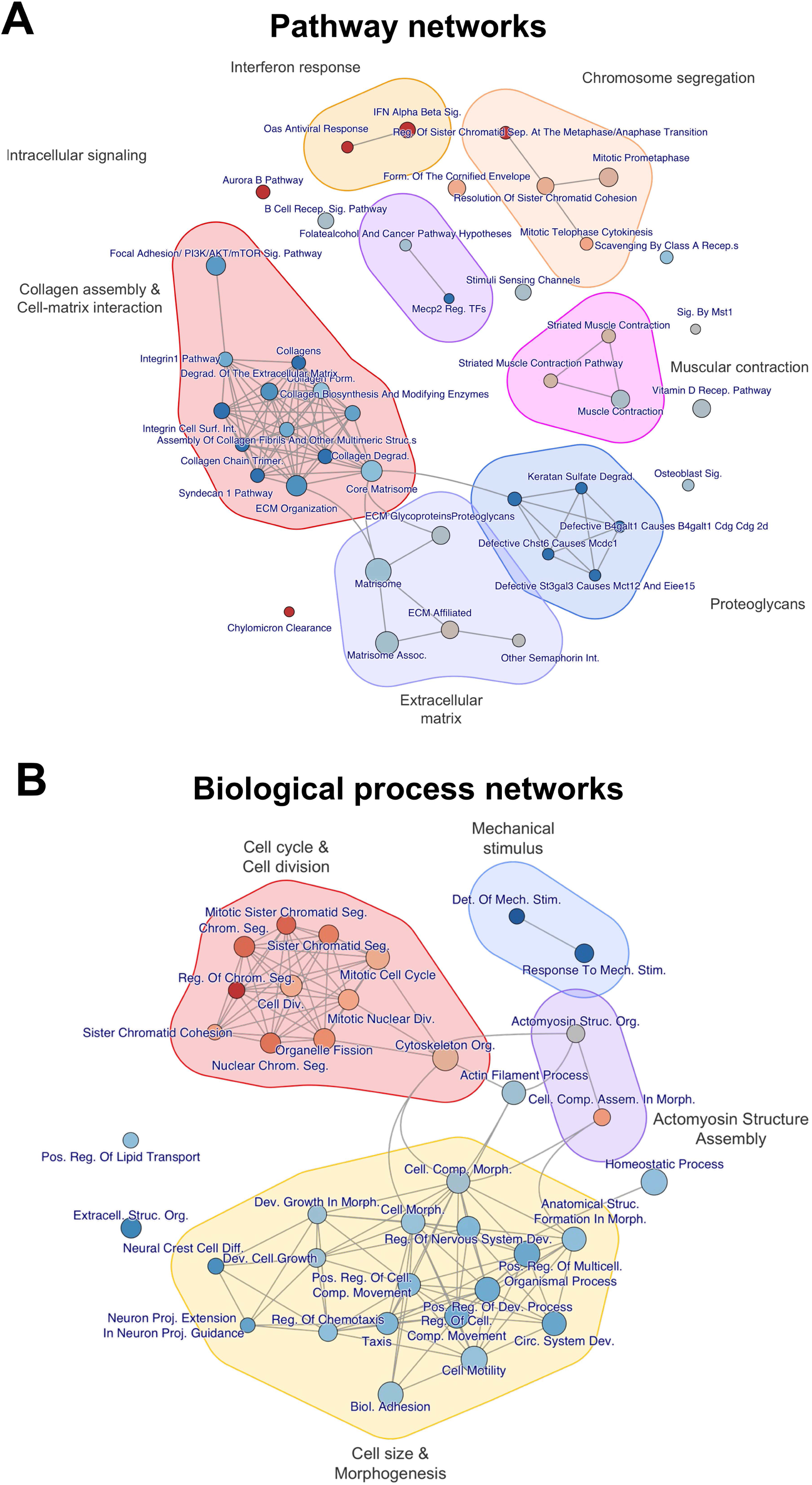
Prediction of PFI-3 molecular mechanism model. **A)** Pathway enrichment network map. Enrichment networks were created using pathways with an adjusted P-value of 0.05, where pathways were colored by direction of gene regulation and were connected between pathways with a Jaccard gene overlap of at least 0.2. **B)** Biological process network map.

Further gene expression analysis showed the downregulation of a subset of 16 genes (Figure 6A), including those crucial for collagen assembly and crosslinking, at both the RNA and protein level (Figure 6B & Figure 6C). For instance, PFI-3 significantly decreased COL11A1, which is involved in collagen assembly [34]. Likewise, Fibronectin, a gene that interacts with collagen, cell receptors, and other ECM components to regulate the collagen synthesis, deposition, and remodeling [35, 36], was also decreased by PFI-3. In addition, we evaluated the activity of the collagen crosslinking enzyme LOX and found a 15% decrease in LOX activity in PFI-3 treated cells on day 3 of reprogramming (Figure 6D). This suggests that PFI-3 may also affect the crosslinking and stabilization of collagen fibers.

**Figure 6.**
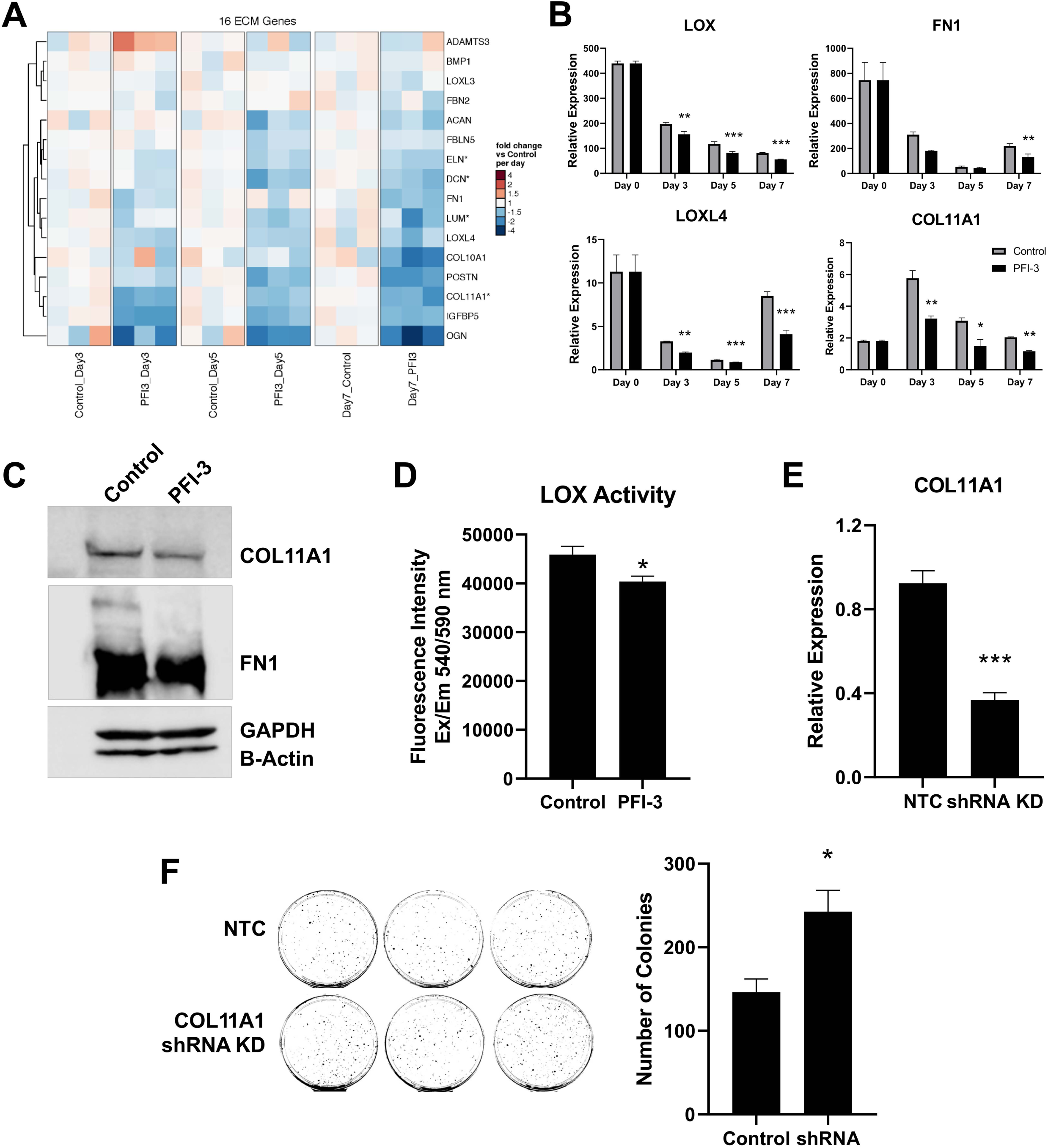
PFI-3 inhibits the expression of genes associated with collagen assembly and crosslinking. **A)** Heatmap of 16 ECM genes which were downregulated by PFI-3**. B)** Quantitative PCR analysis for collagen assembly and collagen crosslinking genes. \**P* value <0.05, n=3. *P* values were 0.012, 0.0006, 0.00013 (LOX), 0.007, 0.0009, 0.0003 (LOXL4), 0.4, 0.06, 0.004 (FN1), 0.002, 0.03, 0.001 (COL11A1). **C)** Protein expression for collagen assembly and collagen crosslinking genes**. D)** LOX enzyme activity assay. \**P* value ≤ 0.05, n=3. *P* values was 0.05. **E)** Quantitative PCR analysis for COL11A1 knockdown efficiency. Human dermal fibroblasts were infected with lentiviruses containing COL11A1 shRNA. Cells were harvested after 72 hours for qPCR analysis. \*\*\**P* value < 0.001, n=3. *P* values was 0.0002. **F)** Alkaline phosphatase (AP) staining of iPS colonies generated with knockdown of COL11A1. Number of AP positive iPS colonies in the PFI-3 treated cells compared with the control cells. \**P* value <0.05, n=3. *P* value was 0.03.

Given the important role of COL11A1 in EMT regulation in cancers [37, 38], we explored the functionality of COL11A1 in reprogramming. We applied shRNA to knockdown COL11A1 in dermal fibroblasts and validated the knockdown efficiency by qRT-PCR (Figure 6E).

Subsequently, these fibroblasts were reprogrammed while keeping COL11A1 suppressed. We found the number of AP-stained iPSC colonies in COL11A1 KD cells were 1.7-fold increase in the COL11A1 KD cells compared to the control (Figure 6F). The result shows that knocking down COL11A1 facilitates iPSC reprogramming and that COL11A1 is not only an EMT regulator but also a reprogramming barrier.

In conclusion, our findings indicate that PFI-3 enhances iPSC reprogramming by promoting MET through repression of genes involved in ECM collagen assembly and collagen crosslinking.

## DISCUSSION

In this study, we evaluated the effect of bromodomain inhibitor PFI-3 on human iPSC reprogramming. Our results indicate that PFI-3 exposure enhances reprogramming efficiency by targeting the gene expression of ECM and promoting the process of MET.

Previous studies have shown that ECM is a barrier to reprogramming [39], with genes associated with ECM organization being repressed during the early stages of the reprogramming process [40]. Our data show that PFI-3 can further repress the gene expression of ECM (Figure 3). Interestingly, while other bromodomain inhibitors have been shown to downregulate fibroblast gene signatures to alter cell identity [41, 42], our study did not observe a significant downregulation of fibroblast markers, such as Vimentin and Thy1. Instead, we found that PFI-3 inhibited EMT master regulator COL11A1 and upregulated MET marker E-cadherin (Figure 4). This suggests that PFI-3 increases cellular plasticity during iPSC reprogramming.

Human dermal fibroblasts have an extensive ECM which is rich in collagen [43]. Fibroblasts are firmly attach to the ECM through the binding of integrins to collagen [44], and integrins link intercellular cytoskeleton to regulate cell adhesion and cell migration [45]. Single-cell imaging reveals that only small and fast dividing cells can be reprogrammed into iPSCs. Because fibroblasts display a stretched, elongated morphology, they have to reduce cell size to achieve a faster proliferation rate at early event of reprogramming [46]. The effect of PFI-3 on ECM gene expression suggests that fibroblasts must overcome the cell-matrix adhesion first to reduce cell size. The downregulation of collagen assembly related genes by PFI-3 could reduce this cell-matrix adhesion and facilitate cell morphology changes.

While the role of ECM in driving EMT induction through mechanotransduction has been extensively studied [47–49], the induction of MET in iPSC reprogramming, particularly from the ECM perspective, remain relatively unexplored. In cancers, collagen crosslinking produces stiffer ECM [20, 21]; integrins then sense this mechanical force and drive rearrangement of actin cytoskeleton to create tension, which triggers a signaling cascade to the nucleus to drive EMT [50, 51]. Our results indicate that PFI-3 modifies the gene expression of ECM, specifically inhibiting collagen assembly and crosslinking, which could lead to a softer ECM. Soft ECM reduces integrin clustering and focal adhesion [52, 53], promoting the expression of E-cadherin and driving polarity establishment and apical-basal junction formation [54]. Studies on the impact of biomaterial stiffness have shown that softer hydrogels led to increased MET and higher reprogramming efficiency in mouse embryonic fibroblasts [55]. Therefore, mechanical cues from the microenvironment not only drive EMT but also induce MET.

Interestingly, our data suggest that MET occurs later in human dermal fibroblast reprogramming, aligning with the activation of pluripotency networks (Figure 4, Figure S3). This result is consistent with previous studies comparing MET timelines between humans and mice [56], highlighting the developmental difference between these species [4]. Our findings imply that ECM remodeling may take longer in humans than in mice due in part to the difference in cell-matrix adhesion. Future experiments comparing ECM gene expression profiles and ECM remodeling between humans and mice will provide further insight. Additionally, we found that PFI-3 enhanced cell immune response during day 7 of reprogramming. This immune response has been shown to improve nuclear reprogramming efficiency in human fibroblasts [57]. The lack of observable upregulation of immune response genes in the early stage of reprogramming suggests that the immune response could be a downstream effect of ECM reorganization. Future experiments measuring ECM stiffness and cross-linked collagen quantification will strengthen these results. It will be also interesting to investigate the effect of PFI-3 on EMT in metastatic cancer cells and protein complexes associated with PFI-3.

In conclusion, our study demonstrates the potential of epigenetic regulators such as the bromodomain inhibitor PFI-3 as tools for enhancing iPSC reprogramming efficiency and highlight the crucial role of ECM changes and cell plasticity in this process. These findings offer compelling insights into the mechanisms behind successful human iPSC reprogramming and open new possibilities for future research in this important therapeutic area.

## ACKNOWLEDGEMENTS

We would like to thank Drs. Elizabeta Gjoneska, Guang Hu, Jackson Hoffman, and Ms. Ginger Muse from NIEHS for critical and thoughtful evaluation of this manuscript.

## AUTHOR CONTRIBUTIONS

Conceptualization, J.Y, T.K.A; Methodology, J.Y, H.K.K., T.K.A.; Formal Analysis, J.Y, H.K.K.; Investigation, J.Y., H.K.K.; Data Curation, J.Y., E. S.; Writing-Original Draft, J.Y., E.S.; Writing-Reviewing Final Draft J.Y., H.K.K., T. K.A., Visualization J.Y, J.M.W.; Supervision, T.K.A; Funding and resources, T.K.A.

## DATA AVAILABILITY

Data are deposited in GEO with the accession number GSE241435.

---

**Supplemental Figure 1.**
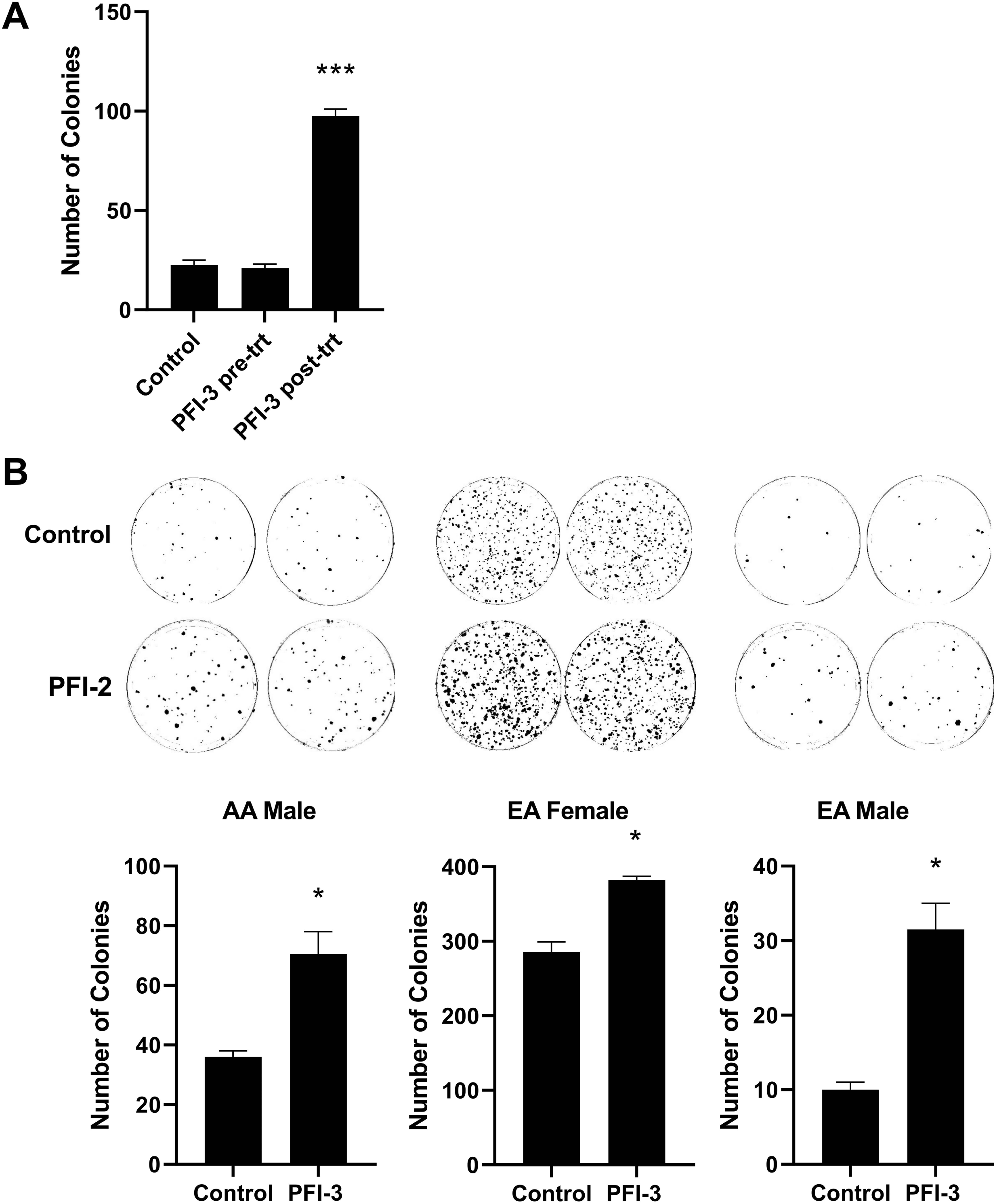
**(A)** PFI-3 pre-treatment has no effect on reprogramming efficiency. **P value <0.001, n=3. P values was 0.0004. **(B)** PFI-3 promotes human iPSC reprogramming among different demographic background. *P value <0.05, n=3. P values were 0.04, 0.02, 0.03.

**Supplemental Figure 2.**
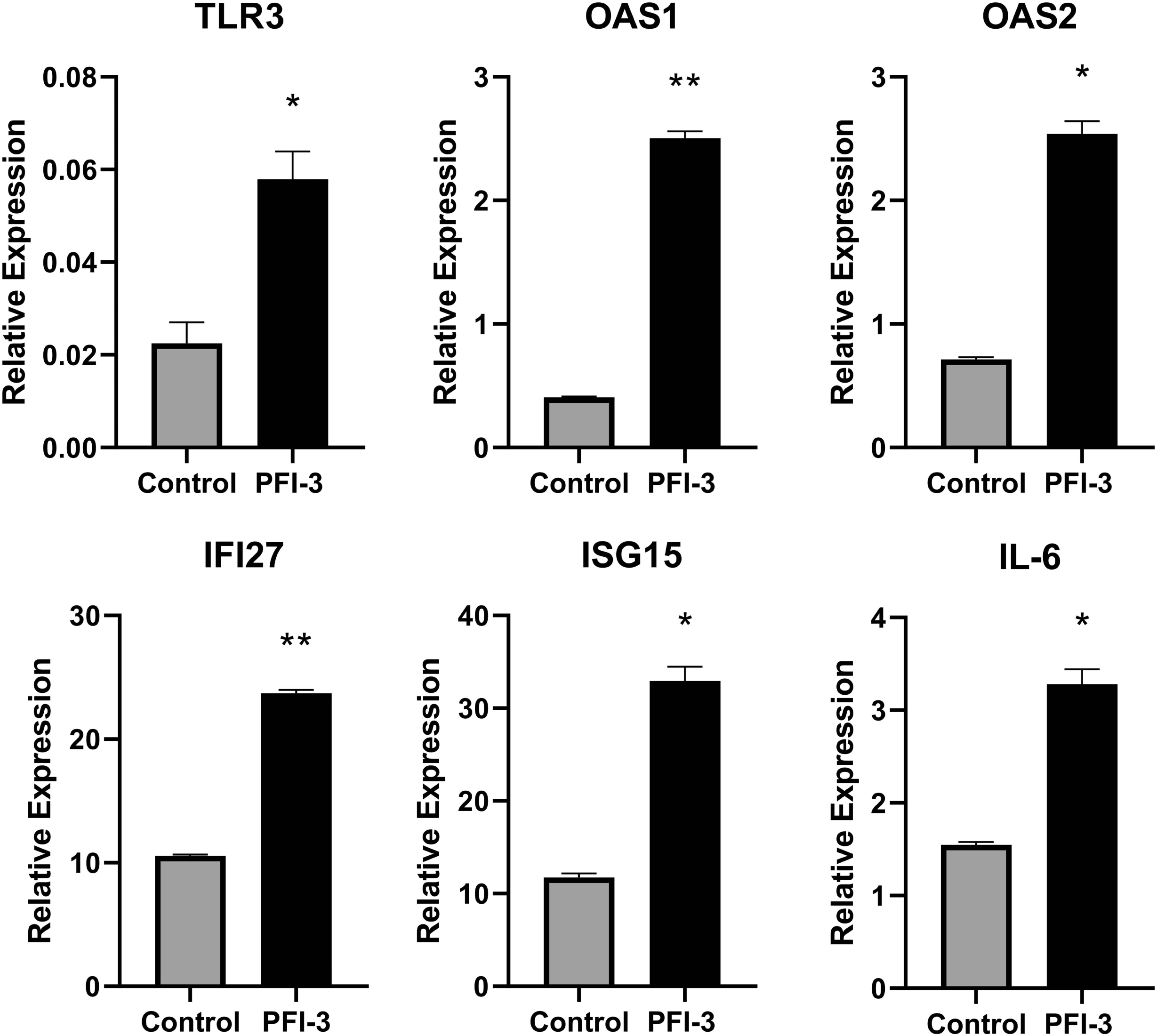
PFI-3 induces innate immune gene expression at day 7 of reprogramming. Quantitative PCR analysis for immune genes TLR3, OAS1, OAS2, IFI27, ISG15, and IL-6 on reprogramming day 7. Transcript level were normalized to TBP expression. *P value <0.05, n=3.

**Supplemental Figure 3.**
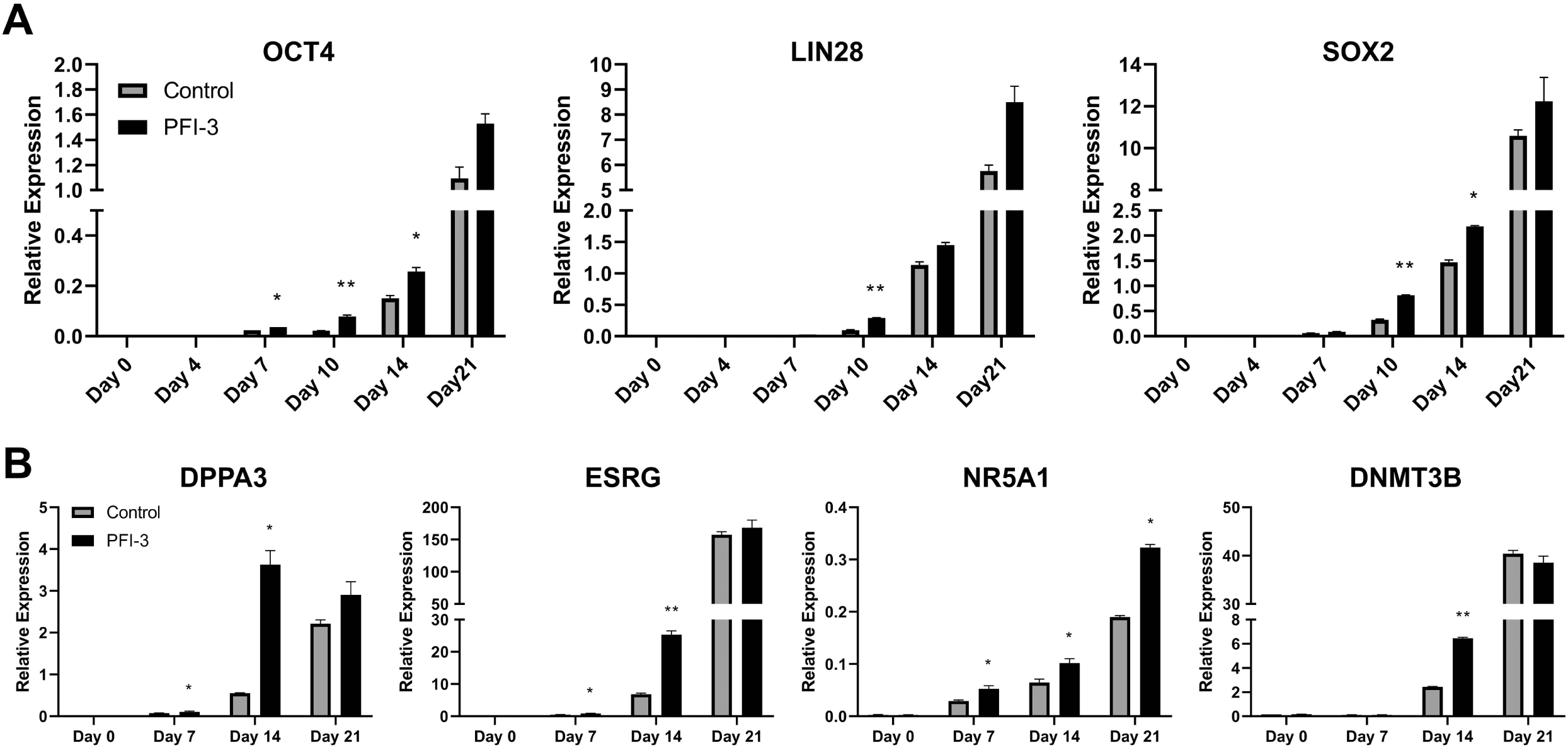
PFI-3 promotes pluripotency network **A)** Quantitative PCR analysis for endogenous pluripotency markers OCT, LIN28, and SOX2 at day 0, 4, 7, 10, 14, and 21 of reprogramming. Transcript level were normalized to TBP expression. *P value <0.05, n=3. **B)** Quantitative PCR analysis for critical reprogramming genes DPPA3, NR5A1, ESRG, and DNMT3B at day 0, 7 and 14. Transcript level were normalized to TBP expression. *P value <0.05, n=3.

